# Geography, not food availability, reflects compositional differences in the bacterial communities associated with larval sea urchins

**DOI:** 10.1101/394486

**Authors:** Tyler J. Carrier, Sam Dupont, Adam M. Reitzel

## Abstract

Determining the principles underlying the assembly, structure, and diversity of symbiont communities remains a focal point of animal-microbiome research. Much of these efforts focus on taxonomic variation within or between animal populations, but rarely test the proportional impacts of ecological components that may affect animal-associated microbiota. Using larvae from the sea urchin *Strongylocentrotus droebachiensis* from the Atlantic and Pacific Oceans, we test the hypothesis that inter-population differences in the composition of animal-associated bacterial communities are more pronounced than intra-population variation due to a heterogeneous feeding environment. Despite significant differences in bacterial community structure within each *S. droebachiensis* larval population (based on food availability, time, development, and phenotype), variation in OTU membership and community composition correlated more strongly with geography. Moreover, across these three locations, 20-30% of OTUs were specific to a single population while less than 10% were shared. Taken together, these results suggest that inter-populational variation in symbiont communities is more pronounced than intra-populational variation. This difference may suggest that ecological variables over broad geographic scales may mask smaller scale ecological variables; however, explicit testing of this potential principle requires common garden experiments as well as molecular and functional manipulations.

## Introduction

Acclimating to environmental variability through morphological, developmental, and/or physiological plasticity is a common trait of animals (Bradshaw, 1965, Boidron-Metairon, 1988, Sterns, 1989, DeWitt *et al.*, 1998, Schlichting & Smith, 2002, West-Eberhard, 2003, Miner *et al.*, 2005). Over the past decade, the appreciation for the role that animal-associated microbiota play in responding to environmental stressors has grown profoundly (Kohl & Carey, 2016, Macke *et al.*, 2016, Shapira, 2016, Theis *et al.*, 2016, Apprill, 2017, Carrier & Reitzel, 2017, Carrier & Reitzel, 2018). When experiencing heterogeneous environments, an animal may recruit, expel, and/or shuffle the relative proportion of associated microbiota (Zilber-Rosenberg & Rosenberg, 2008, Bordenstein & Theis, 2015) to assemble a community with particular molecular pathways for the environmental conditions (Burke *et al.*, 2011, Louca *et al.*, 2016, Roth-Schulze *et al.*, 2018).

Microbial communities associated with animals often vary in response to a multitude of abiotic and biotic factors, including temperature, salinity, diet quality and quantity, season, and habitat-type (see reviews by Kohl & Carey, 2016, Carrier & Reitzel, 2017). Of these, dietary responses are best studied and shown to have a major impact on the composition and function of this community (Kohl & Dearing, 2012, David *et al.*, 2014, Rosenberg & Zilber-Rosenberg, 2016, Sonnenburg *et al.*, 2016). When faced with prolonged diet restriction, the composition and diversity of microbiota associated with both invertebrate and vertebrate hosts shift considerably (Kohl *et al.*, 2014, Carrier & Reitzel, 2018), a response hypothesized to buffer against reduced exogenous nutrients.

Microbial communities associated with animals are also species-specific (*e.g.*, Fraune & Bosch, 2007, Schmitt *et al.*, 2012, Carrier & Reitzel, 2018) and taxonomically variable across the geographical distribution of the host (Dishaw *et al.*, 2014, Marzinelli *et al.*, 2015, Mortzfeld *et al.*, 2015, Marino *et al.*, 2017, Huang *et al.*, 2018). Population-specific microbial communities are partially dependent on environmental conditions (Pantos *et al.*, 2015) and dispersal limitations (Moeller *et al.*, 2017). Despite this taxonomic variation, microbial communities can remain functionally similar due to shared genes across bacterial taxa (Louca *et al.*, 2018, Roth-Schulze *et al.*, 2018). The microbial communities of the bromeliad *Aechmea nudicaulis*, for example, are taxonomically dynamic but functionally consistent between individuals (Louca *et al.*, 2016). Similarly, the bacterial community associated with the green alga *Ulva* spp. are too spatially and temporally variable to define a ‘core’ community; however, nearly 70% of the microbial genes are conserved across the broad geography of *Ulva* spp. (Roth-Schulze *et al.*, 2018).

Efforts to classify the principles underlying the assembly of host-associated microbial communities have primarily been within (*e.g.*, Webster *et al.*, 2011, Webster *et al.*, 2011, David *et al.*, 2014, Lokmer & Wegner, 2015, Carrier & Reitzel, 2018, Carrier *et al.*, 2018) or between (*e.g.*, (Yatsunenko *et al.*, 2012, Dishaw *et al.*, 2014, Marzinelli *et al.*, 2015, Mortzfeld *et al.*, 2015, Marino *et al.*, 2017, Huang *et al.*, 2018) populations (but see Webster *et al.*, 2018). This conceptual discontinuity highlights a gap in understanding the patterns and principles underlying community assembly. To test for differences in assembly and structure, direct comparisons of animal-associated microbial communities within and between populations are needed to determine the relevance of each factor. The fields of sensory, behavioral, and circadian biology have shown that organismal responses to ecological variables are unequal and may be masked based on spatial significance (Beauchard *et al.*, 2017, Hodin *et al.*, 2018, Li *et al.*, 2018, Yang *et al.*, 2018, Kerr & Kemp, In Press). Factors within and between populations, therefore, may have differential impacts on the composition of host-associated microbiota.

Planktotrophic (feeding) larvae are one biological system to compare ecological components and how they influence the composition of animal-associated microbiota. Many planktotrophic larvae (*e.g.*, the pluteus of sea urchins) inhabit heterogeneous nutritional environments and are morphologically and physiologically plastic to food availability (Boidron-Metairon, 1988, Hart & Strathmann, 1994, Miner, 2004, Byrne *et al.*, 2008, Soars *et al.*, 2009, Adams *et al.*, 2011, Miner, 2011, Carrier *et al.*, 2015, McAlister & Miner, 2018). Larvae of the echinoid *Strongylocentrotus droebachiensis*, for example, associate with phenotype-, diet-, and developmental stage-specific bacterial communities (Carrier & Reitzel, 2018, Carrier & Reitzel, 2019). *S. droebachiensis*, in particular, has an Arctic-boreal distribution composed of a multiple ocean population with limited gene flow (Addison & Hart, 2004, Addison & Hart, 2005, Scheibling & Hatcher, 2013) and evidence for potential local adaption (Manier & Palumbi, 2008, Marks *et al.*, 2008).

Using the circumpolar distribution of *S. droebachiensis* and the ability of their larvae to acclimate to heterogeneous feeding environments, we test the hypothesis that inter-population differences in the composition of animal-associated bacterial communities are more pronounced than intra-population variation. To test this, we performed differential feeding experiments on *S. droebachiensis* larvae from populations in the Pacific and Atlantic Oceans (Fig. 1) and used amplicon sequencing to profile the larval-associated bacterial communities.

**Figure 1.**
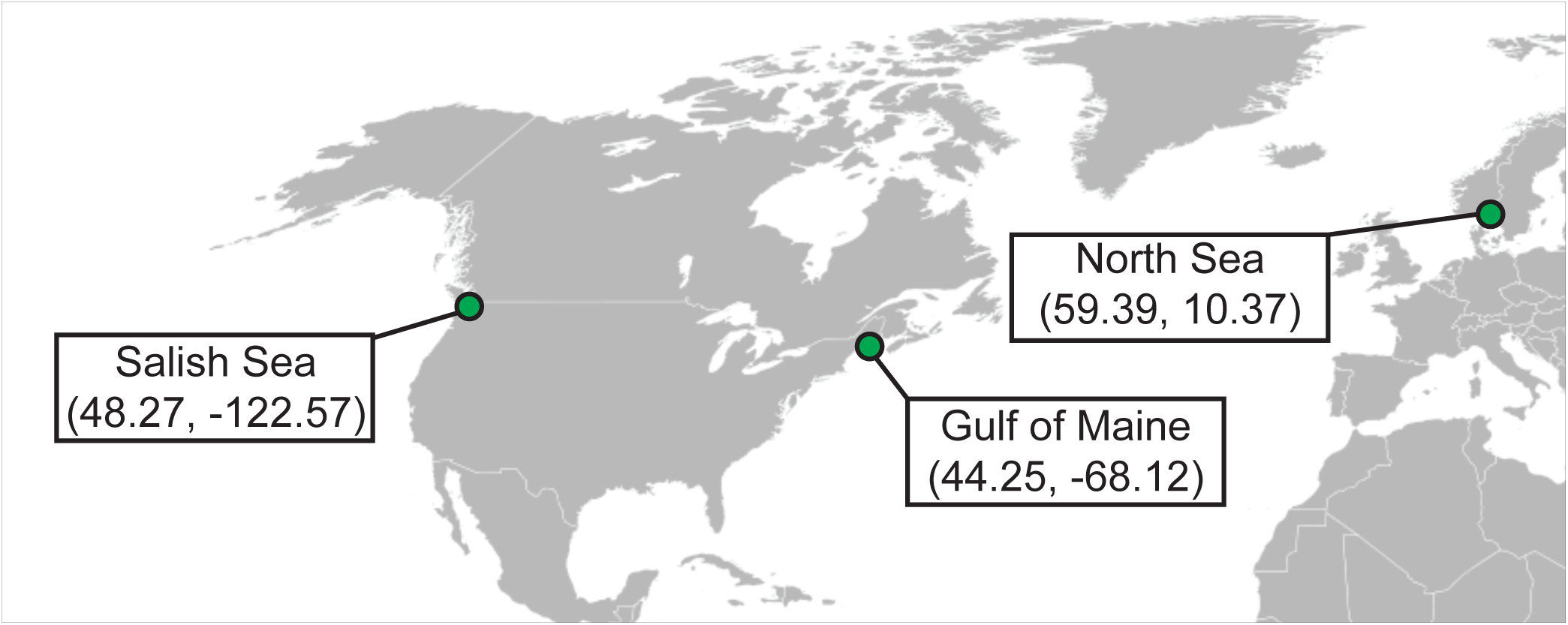
Location of *Strongylocentrotus droebachiensis* experiments. Cartoon representation of adult populations and the geographical distribution for where differential feeding experiments were conducted.

## Materials and Methods

### Adult urchin collection and larval rearing

Adult *S. droebachiensis* were collected from populations in the North Sea in March 2015, the Salish Sea in April 2016, and the Gulf of Maine in February 2017 (Fig. 1). Individuals from the North Sea were collected by divers in Drøbak, Norway (59°39’ N, 10°37’ E) and transported in cold, aerated seawater to the Sven Lovén Centre for Marine Infrastructure (Kristineberg, Sweden). Urchins were maintained in natural seawater and fed *ad libitum* on a live mix of *Ulva lactuca* and *Laminaria* spp. collected from the Kristineberg shoreline. Urchins from the Salish Sea were hand-collected at low tide at Cattle Point, San Juan Island, USA (48°27' N, 122°57' W), transferred to the Friday Harbor Laboratories (FHL; Friday Harbor, WA, USA) within one hour, suspended in sub-tidal cages off the dock at FHL, and fed *Nereocystis* spp. *ad libitum*. Lastly, individuals from the Gulf of Maine were collected by divers in Frenchman Bay, Maine (44°25' N 68°12' W), shipped overnight to the Darling Marine Center (Walpole, ME, USA), and were maintained in flow-through aquaria and fed *Saccharina latissima ad libitum*.

Within two weeks of collections, adult urchins were spawned by a one to two mL intracoelomic injection of 0.50 M potassium chloride. For each population, gametes from three males and females were pooled separately. Fertilization of eggs and larval rearing followed Strathmann (1987), with the exception that embryos and larvae were reared using 5.0-μm filtered seawater (FSW) to include the environmental microbiota. Briefly, embryos were incubated in 1 L of FSW at ambient temperature and salinity. Two hours post-fertilization, seawater was decanted to remove excess sperm. Embryos were then transferred to triplicate 3 L jars and diluted to a density of two individuals•mL^−1^. Beginning at the prism stage, larvae were provided *Rhodomonas lens* at 10,000 cells•mL^−1^, for which was made growth medium-free by centrifugation and resuspension in FSW prior to the introduction to larval cultures. Larval cultures were subsequently diluted to one larva per two mL at advanced developmental stages. Larval cultures were given 90-95% water changes every other day and *R. lens* was replenished at 10,000 cells•mL^−1^.

Monocultures of *R. lens* were grown in f/2 media at room temperature with a combination of ambient and artificial lighting for 24 hours per day (Guillard, 1975).

### Experimental feeding and larval morphometrics

At 48 hours post-fertilization, prism-stage larvae were divided into twelve replicate jars that were sub-divided into four experimental feeding treatments: 10,000, 1,000, 100, or 0 cells•mL^−1^ of *R. lens*. For each experiment, larvae fed 10,000 cells•mL^−1^ were reared through metamorphosis while diet-restricted (1,000 and 100 cells•mL^−1^) and unfed larvae were cultured until developmental stasis (Table S1). Larvae (n=100) from each replicate for each treatment were sampled weekly. Immediately after sampling, larval samples were concentrated into a pellet using a microcentrifuge, the FSW was removed with a sterile glass pipette, and pelleted larvae were then preserved in RNAlater (Thermo Scientific, Massachusetts, USA) and stored at −20°C before DNA extractions.

Complementary to sampling *S. droebachiensis* larvae, the environmental microbiota from the seawater was also sampled. When larval cultures were sampled, triplicate ∼1-L of 5.0-μm FSW was filtered onto a 0.22-µm Millipore filter to retain the environmental microbiota. Full filter disks were then preserved in RNAlater and stored at −20 °C before DNA extractions.

In addition to sampling larvae to assay the associated bacterial communities, twenty larvae (n=20) from a single replicate of each dietary treatment were sampled for morphometric analysis. Larvae were imaged using a compound microscope (Salish Sea: Nikon Eclipse E600; camera: QImaging MicroPublisher 5.0 RTV; Gulf of Maine: Zeiss Primo Star HD digital microscope; North Sea: Leica stereomicroscope) and morphometrics (length of larval body, post-oral arms, and stomach area; Figs. 1, S1-2) were performed using ImageJ (v. 1.9.2; Schneider *et al.*, 2012). Statistical differences in larval size and stomach volume were compared using a two-way analysis of variance (ANOVA; JMP Pro v. 13) to test for differences over time and across feeding conditions. Moreover, differences between sites were compared with a one-way ANOVA. Where statistical differences were observed (p<0.05), a Tukey’s post-hoc test were used to determine the affect at each time point and for each diet.

### Assaying microbial communities

Total DNA was extracted from larval samples using the GeneJet Genomic DNA Purification Kit (Thermo Scientific, Massachusetts, USA). For FSW samples, eDNA was extracted using the FastDNA Spin Kit for Soil (MP Biomedical, Illkirch, France). DNA was then quantified using the NanoDrop 2000 UV-Vis Spectrophotometer (Thermo Scientific, Massachusetts, USA) and diluted to 5 ng•μL^−1^ using RNase/DNase-free water.

Bacterial sequences were amplified using universal primers for the V3/V4 regions of the 16S rRNA gene (Forward: 5’ CTACGGGNGGCWGCAG, Reverse: 5’ GACTACHVGGGTATCTAATCC; Klindworth *et al.*, 2013). Products were purified using the Axygen AxyPrep Mag PCR Clean-up Kit (Axygen Scientific, New York, USA), indexed via PCR using the Nextera XT Index Kit V2 (Illumina, California, USA), and then purified again. At each of these three steps, fluorometric quantitation was performed using a Qubit (Life Technologies, California, USA) and libraries were validated using a Bioanalyzer High Sensitivity DNA chip (Agilent Technologies, California, USA). Illumina MiSeq sequencing (v3, 2×300 bp paired-end reads) was performed at the University of North Carolina at Charlotte. See, Table S2 for PCR recipe and thermal profiles.

Forward and reverse sequences (that are freely available on the Dryad Digital Repository) were paired and trimmed using PEAR (Zhang *et al.*, 2014) and Trimmomatic (Bolger *et al.*, 2014), respectively, and converted from fastq to fasta using custom script (see, Note S1). Chimeric sequences were then detected using USEARCH (Edgar *et al.*, 2011) and removed using filter_fasta.py prior to analysis of bacterial 16S rRNA sequences. Using QIIME 1.9.1 (Caporaso *et al.*, 2010) and Greengenes (v. 13.5), bacterial 16S rRNA sequences were analyzed and grouped into operational taxonomic units (OTUs) based on a minimum 97% similarity. The biom table generated by pick_open_reference_otus.py was filtered of OTUs with ten or less sequences as well as sequences matching the cryptophytes (*i.e.*, *R. lens*; as per Carrier & Reitzel, 2018).

Using the filtered biom table and “biom summarize-table” to count total sequences per sample, the rarefaction depth of 3,193 reads was determined and applied to all subsequent analyses (Fig. S3). Alpha diversity (*i.e.*, Fisher’s alpha, Shannon equitability, Faith’s phylogenetic distance, and observed OTUs) was calculated using alpha_diversity.py and compared statistically using one-way ANOVAs in JMP. Beta diversity was calculated using the unweighted and weighted UniFrac (Lozupone & Knight, 2005) as part of jackknifed_beta_diversity.py. These values were compared using principal coordinate analyses (PCoA), recreated using make_2d_plots.py, and stylized for presentation in Adobe Illustrator CS6. Community similarity across phenotypes, dietary states, developmental stages, and biogeography were compared statistically using an analysis of similarity (ANOSIM), permutational multivariate analysis of variance using distance matrices (ADONIS), and permutational multivariate analysis of variance (PERMANOVA) using compare_categories.py. Shared OTUs between urchin populations were determined using compute_core_microbiome.py and shared_phylotypes.py. Average bacterial community for each urchin population as well as across populations and the shared community was generated using summarize_taxa_through_plots.py, visualized with Prism 7 (GraphPad Software), and stylized for presentation in Adobe Illustrator CS6.

A step-by-step listing of QIIME scripts used to convert raw reads to OTUs for visualization of these data is located in Note S1.

## Results

### Morphological plasticity of larval sea urchin

Diet-induced morphological plasticity was observed for *S. droebachiensis* larvae from each population (Figs. S1-2; Tables S3-4), with the extent and direction of expression being location-specific (ANOVA, p<0.0001; Fig. 2). Diet-restricted larvae from the Salish Sea and Gulf of Maine, in general, exhibited a higher post-oral arm to mid-body line ratio than *ad libitum* counterparts (ANOVA, p<0.0001; Figs. S1-2; Tables S3-4), even though analyses of Gulf of Maine larvae were partially confounded by developmental stage (Table S1). Larvae from the North Sea, however, exhibited the opposite response: *ad libitum* feeding induced a higher post-oral arm to mid-body line ratio than diet-restricted individuals (ANOVA, p<0.0001; Fig. S1-2; Tables S3-4).

**Figure 2.**
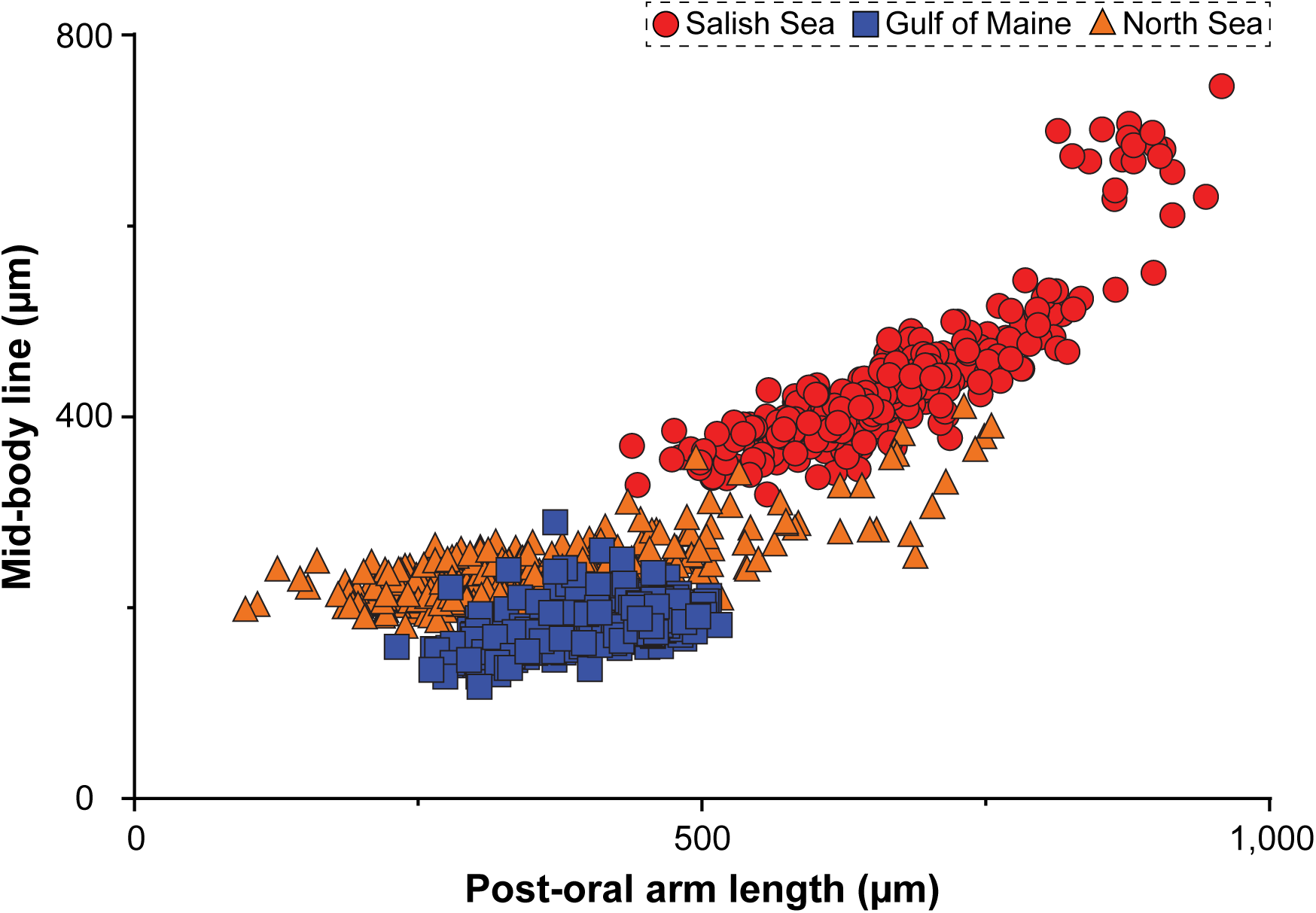
Morphology of *Strongylocentrotus droebachiensis* larvae from three populations. Post-oral arm and mid-body line lengths for *S. droebachiensis* larvae from the Salish Sea, Gulf of Maine, and North Sea, with feeding quantity, developmental stage, and time being collapsed. (See, Figs. S1-2 for the sub-division of larval features.)

### Structural comparisons of bacterial communities within populations

*S. droebachiensis* larvae from each population associated with a diet-specific bacterial community (Figs. 3A-C, S4-7; Table S6). Larvae from the Salish Sea and Gulf of Maine exhibited similar diet-specific community-level patterns in both alpha- and beta-diversity (Figs. 3A-B, S4-6; Table S6), where the bacterial community associated with food restricted larvae were more similar to each other than to well-fed counterparts. Larvae from the North Sea, on the other hand, exhibited the opposite response (Figs. 3C, S4, S7; Table S6), where diet-specific bacterial communities were observed (Table S6) except that all food concentrations were more similar to each other than to starved larvae (Fig. S7; Table S6).

**Figure 3.**
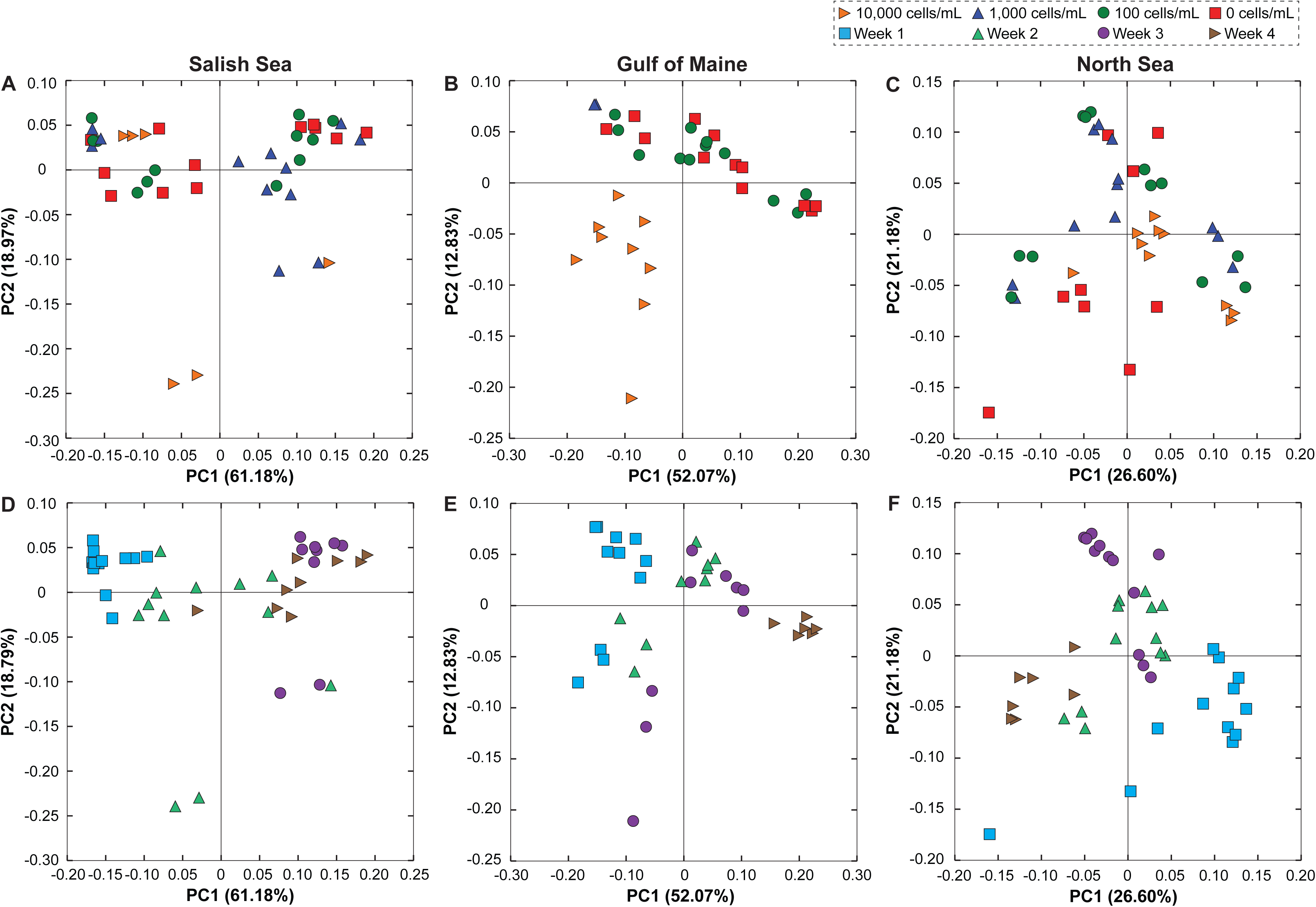
Dietary and temporal shifts in the bacterial communities associated with *Strongylocentrotus droebachiensis* larvae. Community similarity based on *S. droebachiensis* larval-associated bacterial communities based on food availability (A-C) over a multi-week exposure (D-F) for populations from the Salish Sea (A, D), Gulf of Maine (B, E), and North Sea (C, F). Comparisons between food availability (A-C) and over time (D-F) are based on weighted UniFrac comparisons.

The bacterial communities associated with larvae from each sea urchin population also varied temporally in both alpha- and beta-diversity (Figs. 3D-F, S4, S8-10; Table S6). Temporal changes in community diversity (*i.e.*, alpha-diversity) were reflective of culturing environment: larval populations from the Atlantic Ocean increased in weeks one through three and declined in week four while the larval population from the Pacific Ocean decreased in the first three weeks and increased in week four (Table S5). Compositional changes (*i.e.*, beta-diversity) in the associated bacterial communities for each larval population, on the other hand, experienced similar time-dependent shifts from week one though four (Figs. 3D-F, S4).

Complementary to dietary and temporal shifts in community structure and composition, larval-associated bacterial communities for *S. droebachiensis* varied significantly with larval morphology (Fig. S11; Table S7) and developmental stage (Figs. S8-10; Table S7). Variation in the larval-associated bacterial communities across food availability, time, morphology, and development, however, appeared to not co-depend on the bacterial taxa observed in the seawater, as the composition of larval-associated and environmental microbiota were significantly different (Fig. S12; Table S7).

### Differences in the bacterial communities of urchin populations

Despite differences in bacterial community structure within each *S. droebachiensis* larval population, variation in OTU membership and composition correlated best with geography (ANOSIM, unweighted UniFrac, p<0.001; ANOSIM, weighted UniFrac, p<0.001; Figs. 4A-B, S13-15; Table S8). Community relatedness of larval-associated microbiota in the Western and Eastern Atlantic Ocean was more similar to each other than to the Pacific Ocean population (Fig. 4 A-B). Moreover, bacterial community diversity and structure were significantly different between these three populations (Fisher’s alpha: F-ratio = 142.2, *p*<0.001; Shannon equitability: F-ratio = 110.1, *p*<0.001; Faith’s phylogenetic distance: F-ratio = 159.2, *p*<0.001; observed OTUs: F-ratio = 124.2, *p*<0.001; Fig. 4C; Table S8). For each of the tested indices, bacterial communities of Gulf of Maine larvae were taxonomically richest and most diverse while Salish Sea larvae were the least rich and diverse, leaving North Sea larvae as intermediate (Tukey’s test, *p*<0.05 for each; Fig. 4C; Table S8).

**Figure 4.**
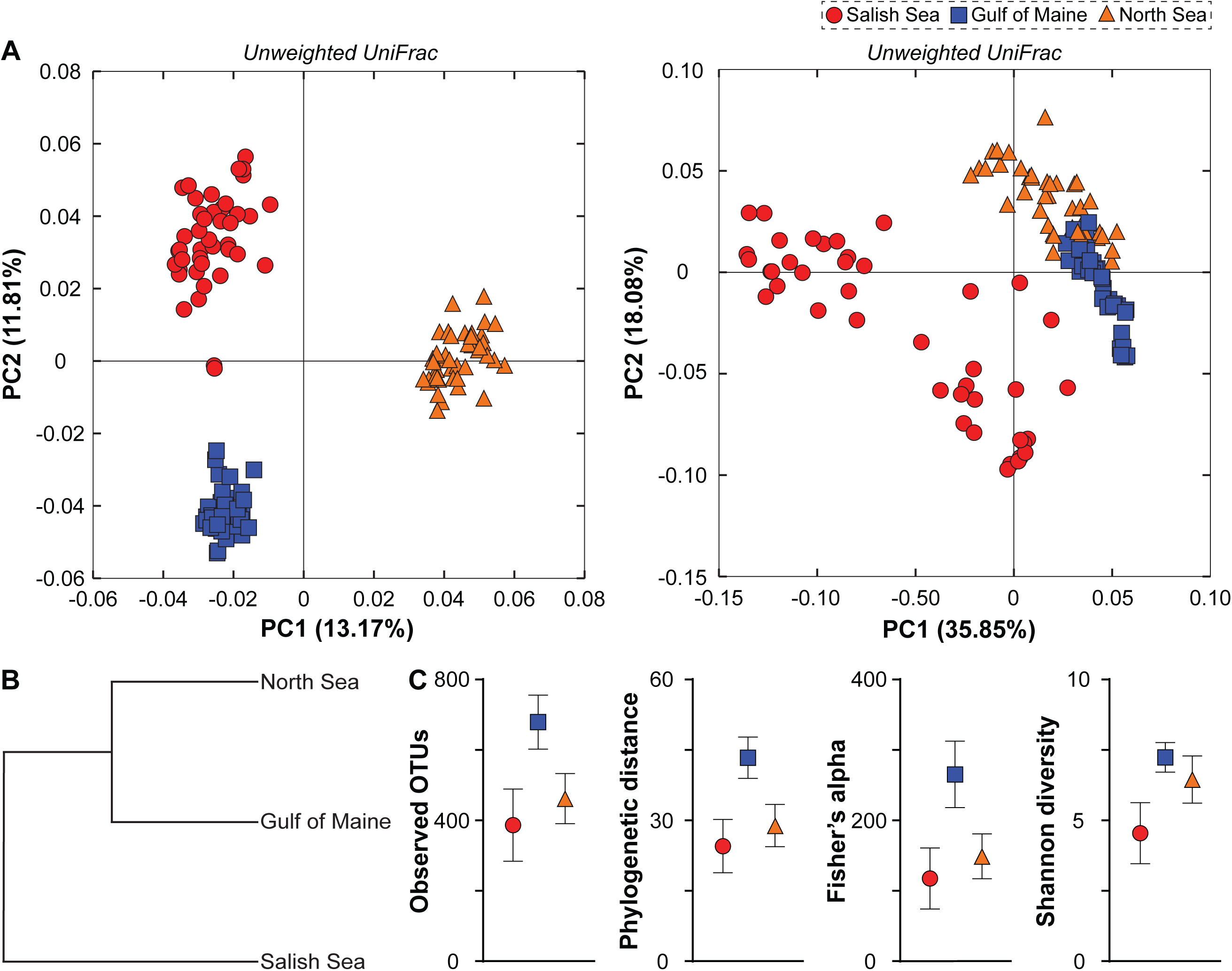
Structure of the bacterial communities associated with *Strongylocentrotus droebachiensis* larvae from three populations. Community similarity of *S. droebachiensis* larval-associated bacterial communities between three geographic locations based on (A) unweighted and weighted UniFrac comparisons, with the relatedness of the weighted UniFrac represented by a microbial dendrogram (B). Diversity within each population was estimated using four (C) alpha metrics: observed OTUs, Faith’s phylogenetic distance, Fisher’s alpha, and Shannon diversity.

*S. droebachiensis* larvae from these populations primarily associated with seven bacterial classes from three phyla: Saprospirae and Flavobacteriia (Bacteroidetes), Alpha-, Beta-, Delta-, and Gammaproteobacteria (Proteobacteria), and Verrucomicrobiae (Verrucomicrobia) (Fig. 5A). Of these classes, Flavobacteriia, Alpha- and Gammaproteobacteria represented at least 10% of the relative community for each population, where Gammaproteobacteria was the most abundant clade at 20-50% (Fig. 5A). Of the hundreds of OTUs associated with *S. droebachiensis* larvae (Fig. S1), less than 10% were shared across populations, with 20-30% being specific to a single population and 3-11% being shared between two populations (Fig. 5B). This shared community was similar to the full communities for each population, in that it was composed primarily by Flavobacteriia (24.7%), Alpha-(28.1%) and Gammaproteobacteria (34.9%) (Fig. 5A).

**Figure 5.**
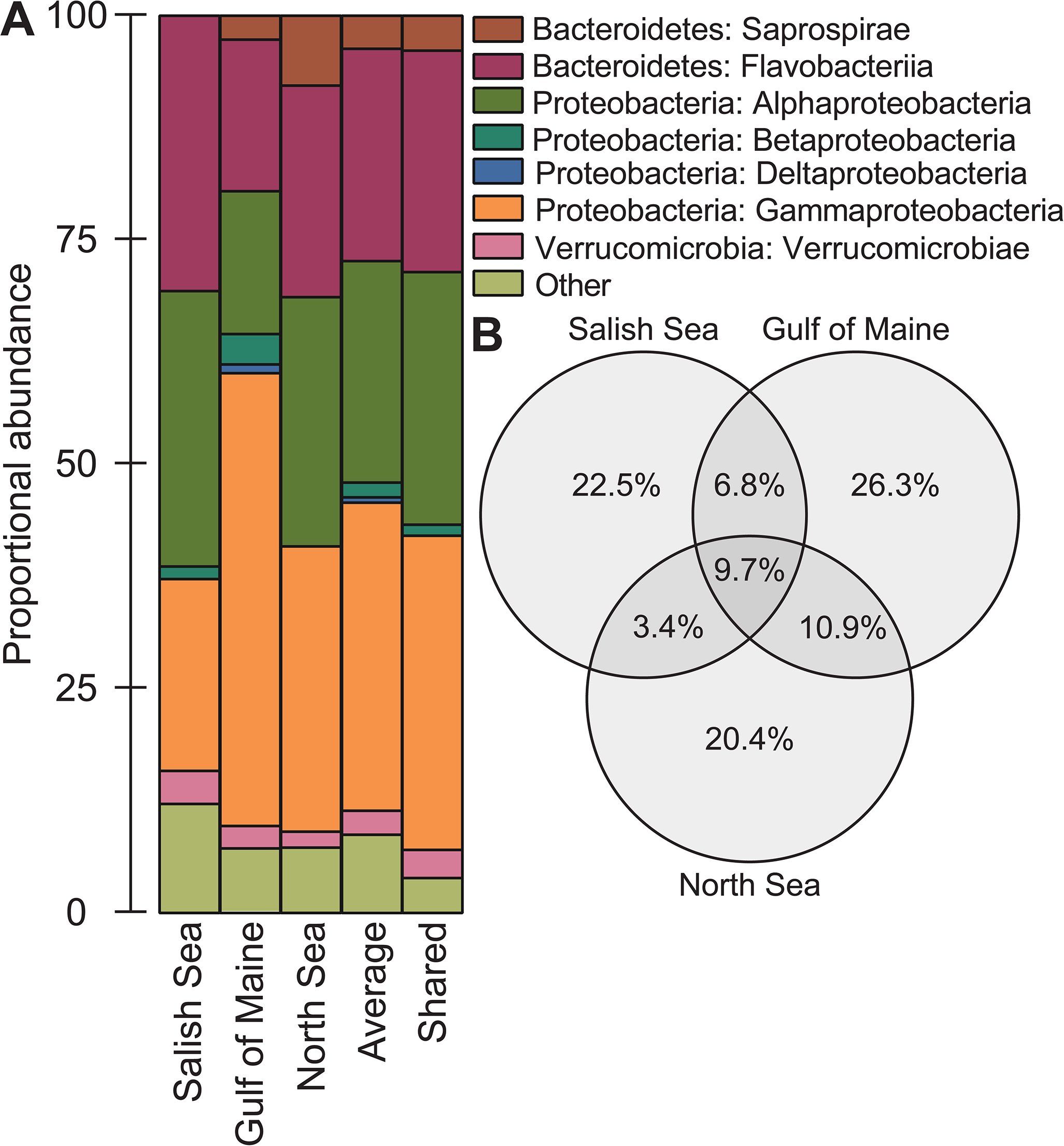
Bacterial taxa associated with *Strongylocentrotus droebachiensis* larvae. (A) Class-level profiles of the bacterial communities associated with *S. droebachiensis* larvae from the Salish Sea, Gulf of Maine, and North Sea as well as the mean community across populations and the shared taxa between each population. (B) Distribution of OTUs associated with *S. droebachiensis* larvae uniquely found in each population, common between two populations, and shared between all populations.

## Discussion

Principles underlying the assembly and diversity of host-associated microbial communities has remained a focal point of host-microbiome research over the last few decades (Ley *et al.*, 2006, Ley *et al.*, 2008, Zilber-Rosenberg & Rosenberg, 2008, Rosenberg *et al.*, 2009, Bright & Bulgheresi, 2010, Bordenstein & Theis, 2015, Coyte *et al.*, 2015, Gilbert *et al.*, 2015, Marzinelli *et al.*, 2015, Brooks *et al.*, 2016, Macke *et al.*, 2016, Shapira, 2016, Carrier & Reitzel, 2017, Huang *et al.*, 2018, Miller *et al.*, 2018). Efforts to partition and categorize variation in community composition have suggested that evolutionary history (Ley *et al.*, 2006, Ley *et al.*, 2008, Zilber-Rosenberg & Rosenberg, 2008, Brooks *et al.*, 2016, Shapira, 2016), physiology (Kohl *et al.*, 2016, Kohl & Carey, 2016, Buckley & Rast, 2017), and environmental conditions (Alberdi *et al.*, 2016, Kohl & Carey, 2016, Carrier & Reitzel, 2017, Miller *et al.*, 2018) primarily drive these differences. The majority of this previous work, however, focuses on two spatiotemporal scales: within (Webster *et al.*, 2011, Webster *et al.*, 2011, David *et al.*, 2014, Lokmer & Wegner, 2015, Carrier & Reitzel, 2018, Carrier *et al.*, 2018) or between (Dishaw *et al.*, 2014, Marzinelli *et al.*, 2015, Mortzfeld *et al.*, 2015, Marino *et al.*, 2017, Huang *et al.*, 2018) host populations (but see Webster *et al.*, 2018). The potential to distinguish and discern intra- and inter-population processes for a single species highlights a potential conceptual gap in understanding the patterns and principles underlying microbiome assembly.

Comparing the bacterial communities associated with *S. droebachiensis* larvae across feeding environments and geographical location supports two primary findings. First, larvae from each population associate with bacterial communities correlated with food availability, time, development, and phenotype. Second, despite variation in the microbiota within each population of urchin larvae, the composition of *S. droebachiensis* larval-associated bacterial communities best correlated with geography.

Larval marine invertebrates associate with microbial communities that are diverse and dynamic yet specific to host species and distinct from the environmental microbiota (Webster *et al.*, 2011, Carrier *et al.*, 2018, Carrier & Reitzel, 2018, Carrier *et al.*, 2018, Carrier & Reitzel, 2019). The bacterial community associated with larval echinoderms, in particular, shifts with food availability (Carrier & Reitzel, 2018, Carrier *et al.*, 2018), embryonic and larval development (Carrier & Reitzel, 2018, Carrier & Reitzel, 2019), phenotype (Carrier & Reitzel, 2018), disease (Carrier *et al.*, 2018), and physiology (Buckley & Rast, 2017). Common approaches for investigating larval-microbe interactions involve single location studies of a widely distributed taxa (see Webster *et al.*, 2011, Carrier & Reitzel, 2018, Carrier *et al.*, 2018, Carrier & Reitzel, 2019). The data presented here suggest that community-level shifts in response to biotic (development and phenotype) and abiotic (food availability and time) factors occur at multiple locations across a population with limited gene flow. Three widely-distributed sea urchin species (including *S. droebachiensis*) have previously been shown to associate with food-elicited phenotype-specific bacterial communities (Carrier & Reitzel, 2018). It may, therefore, be hypothesized that phenotype-specific microbiota for these and, potentially, other species of larval echinoderms (*e.g.*, *Acanthaster* sp. in the Pacific Ocean, including on the Great Barrier Reef; Wolfe *et al.*, 2015, Wolfe *et al.*, 2015, Wolfe *et al.*, 2017, Carrier *et al.*, 2018) occur throughout the geographical distribution of the species.

Marine invertebrate larvae are not unique in their ability to respond to environmental variability by exhibiting shifts in the associated bacterial community (Kohl & Carey, 2016, Carrier & Reitzel, 2017). Several groups of marine and terrestrial vertebrates (*e.g.*, mammals, reptiles, and fish) respond to diet variability and abiotic stress by restructuring their gut microbiota that, in turn, are likely to differ in community function (Muegge *et al.*, 2011, Kohl & Dearing, 2012, David *et al.*, 2014, Schmidt *et al.*, 2015, Kohl *et al.*, 2016, Kohl & Carey, 2016, Kohl *et al.*, 2018). Similar to larval marine invertebrates, marine and terrestrial vertebrates can be widely-distributed and, thus, multiple genetically distinct populations may exhibit similar community-levels shifts in microbiota at multiple locations across these populations. To our knowledge, however, parallel responses between populations for vertebrates and other taxa have not yet been documented.

Inter-populational variation in the composition of *S. droebachiensis* larval-associated bacterial communities was more pronounced than intra-populational differences. While the mechanisms driving this result is unknown, genetic and ecological differences between *S. droebachiensis* populations are likely contributing factors (Pantos *et al.*, 2015, Moeller *et al.*, 2017). *S. droebachiensis* from the Salish Sea, Gulf of Maine, and North Sea have some degree to genetic isolation (Addison & Hart, 2004, Addison & Hart, 2005, Cowen & Sponaugle, 2009, Norderhaug *et al.*, 2016). Moreover, each urchin population inhabit environments that differ in physical (Otto *et al.*, 1990, Pettigrew *et al.*, 2005, Sutherland & MacCready, 2011) and biological (Winter *et al.*, 1975, Thomas *et al.*, 2003, McQuatters-Gollop *et al.*, 2007) factors. Inter-populational variation in the bacterial communities associated with *S. droebachiensis* larvae may, therefore, have been due to differences in host genetics and the oceanographic environment, a result that requires future explicit testing using common garden experiments and molecular manipulations (Williams & Carrier, 2018).

Population-specific bacterial communities for *S. droebachiensis* larvae are consistent with the hypothesis that host-associated microbial communities are taxonomically variable across host geography (Dishaw *et al.*, 2014, Marzinelli *et al.*, 2015, Mortzfeld *et al.*, 2015, Marino *et al.*, 2017, Huang *et al.*, 2018). Studies using diverse animal taxa (*e.g.*, sponges, corals, fish, and humans) support that the environment, not genotype, principally drives community differences (Sullam *et al.*, 2012, Burgsdorf *et al.*, 2014, Pantos *et al.*, 2015, Rothschild *et al.*, 2018). Genetic comparison of *S. droebachiensis* indicates that individuals from the Salish Sea and Gulf of Maine sub-populations are more related to each other than to the North Sea sub-population (Addison & Hart, 2004, Addison & Hart, 2005, Scheibling & Hatcher, 2013). Microbial communities from these populations, however, suggest the contrary: larvae from the Gulf of Maine and North Sea are more similar to each other than to Salish Sea larvae. This result reinforces the hypothesis that environment (or, here, geography) is a larger influence on community assembly than genetic similarity. The mechanistic linkage between host genetics, the environment, and microbial community, however, is unknown.

By comparing host-associated bacterial communities across feeding environments and biogeography in larval sea urchins, we propose that inter-populational variation in symbiont communities is more biologically significant than intra-populational variation. This difference suggests that larger scale ecological variables that correspond to different geographic locations may mask smaller scale ecological variables (*e.g.*, feeding, time, development, and phenotype). Explicit testing with diverse animal taxa by experimental manipulations and functional characterization are merited to understand the priority of exogenous factors in the composition of host-associated microbial communities.

## Acknowledgements

We thank the staffs of the Friday Harbor Laboratories, Darling Marine Center, and Sven Lovén Centre for Marine Infrastructure for facility access and logistical assistance; Jason Hodin and Billie Swalla for providing laboratory space; Richard Strathmann for collection assistance; Daniel Janies for sequencing resources; Karen Lopez for technical assistance with sequencing; and Jason Macrander for bioinformatics assistance.

T.J.C. was supported by an NSF Graduate Research Fellowship, Charles Lambert Memorial Endowment fellowship from the Friday Harbor Laboratories, and a Sigma Xi Grants-in-Aid of Research grant; S.D. was funded by the Centre for Marine Evolutionary Biology and supported by a Linnaeus grant from the Swedish Research Councils VR and Formas; and A.M.R. was supported by Human Frontier Science Program Award RGY0079/2016.

---

**Figure S1.**
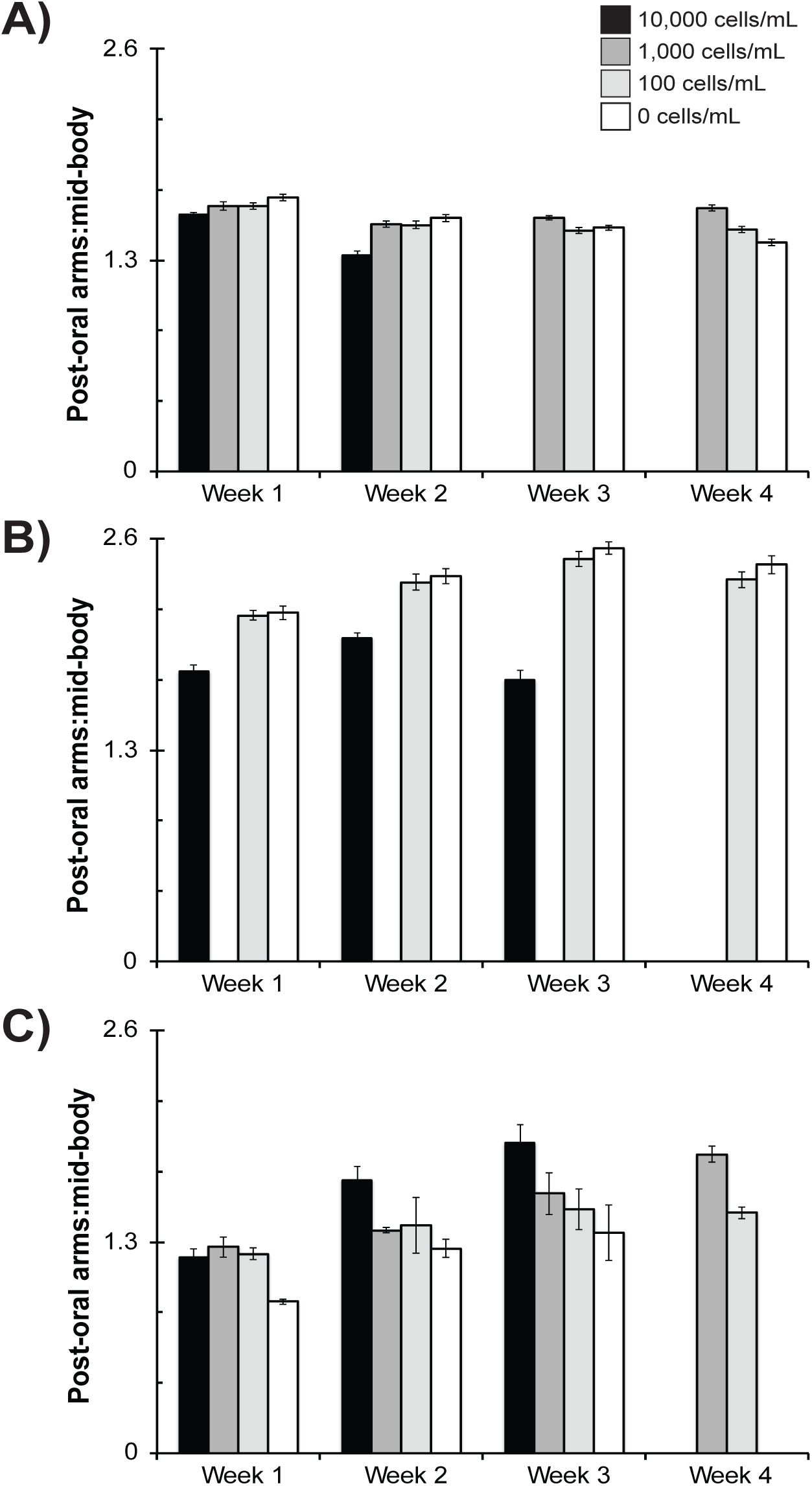
*Strongylocentrotus droebachiensis* larvae alter phenotype to feeding environment. Ratio between the post-oral arm and mid body line (mean ± standard error; n=20) for *S. droebachiensis* from the Salish Sea (A), Gulf of Maine (B), and North Sea (C) larvae having been fed either 10,000 (black), 1,000 (dark grey), 100 (grey), and 0 cells•mL^−1^ (white).

**Figure S2.**
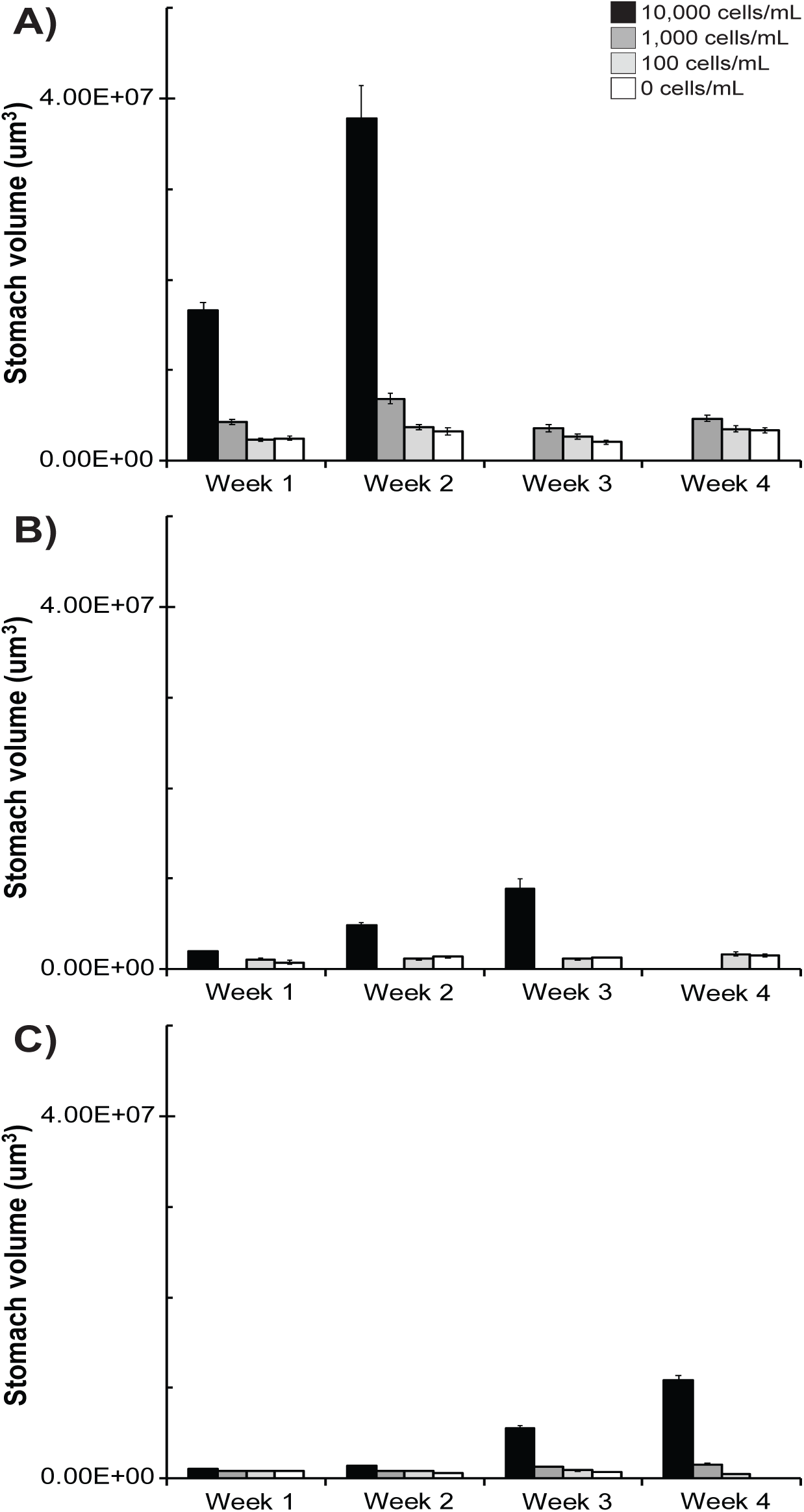
*Strongylocentrotus droebachiensis* larvae alter gut volume to feeding environment. Gut volume (mean ± standard error; n=20) for larval *S. droebachiensis* from the Salish Sea (A), Gulf of Maine (B), and North Sea (C) having been fed either 10,000 (black), 1,000 (dark grey), 100 (grey), and 0 cells•mL^−1^ (white).

**Figure S3.**
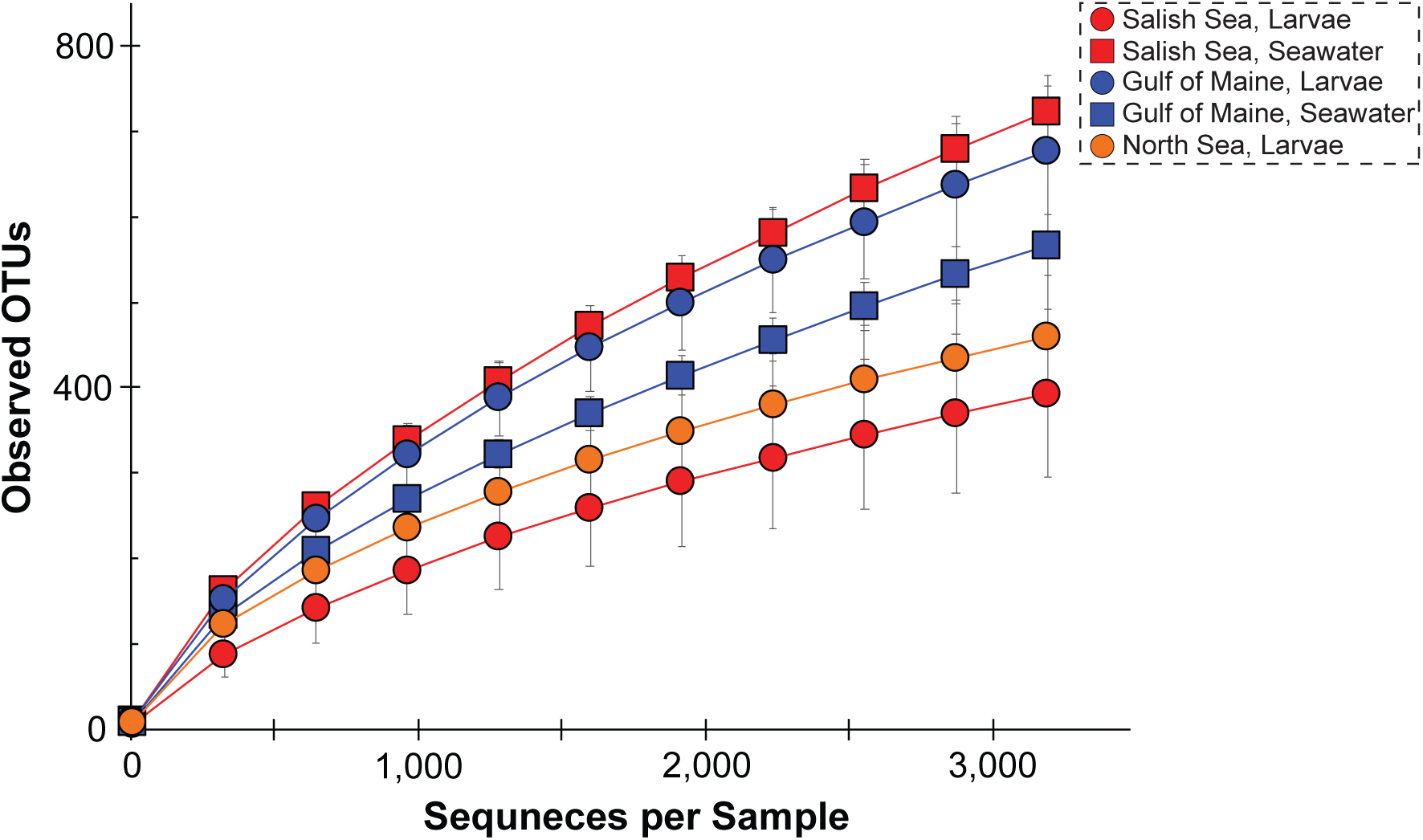
Alpha rarefaction curves for *Strongylocentrotus droebachiensis* larvae from three geographic locations and of the seawater. Alpha rarefaction curves for the associated microbiota for *S. droebachiensis* larvae (circles) from the Salish Sea (blue), Gulf of Maine (red), and North Sea (yellow) and seawater (squares) based on the rarefaction depth (3,193 sequences) used for all analyses.

**Figure S4.**
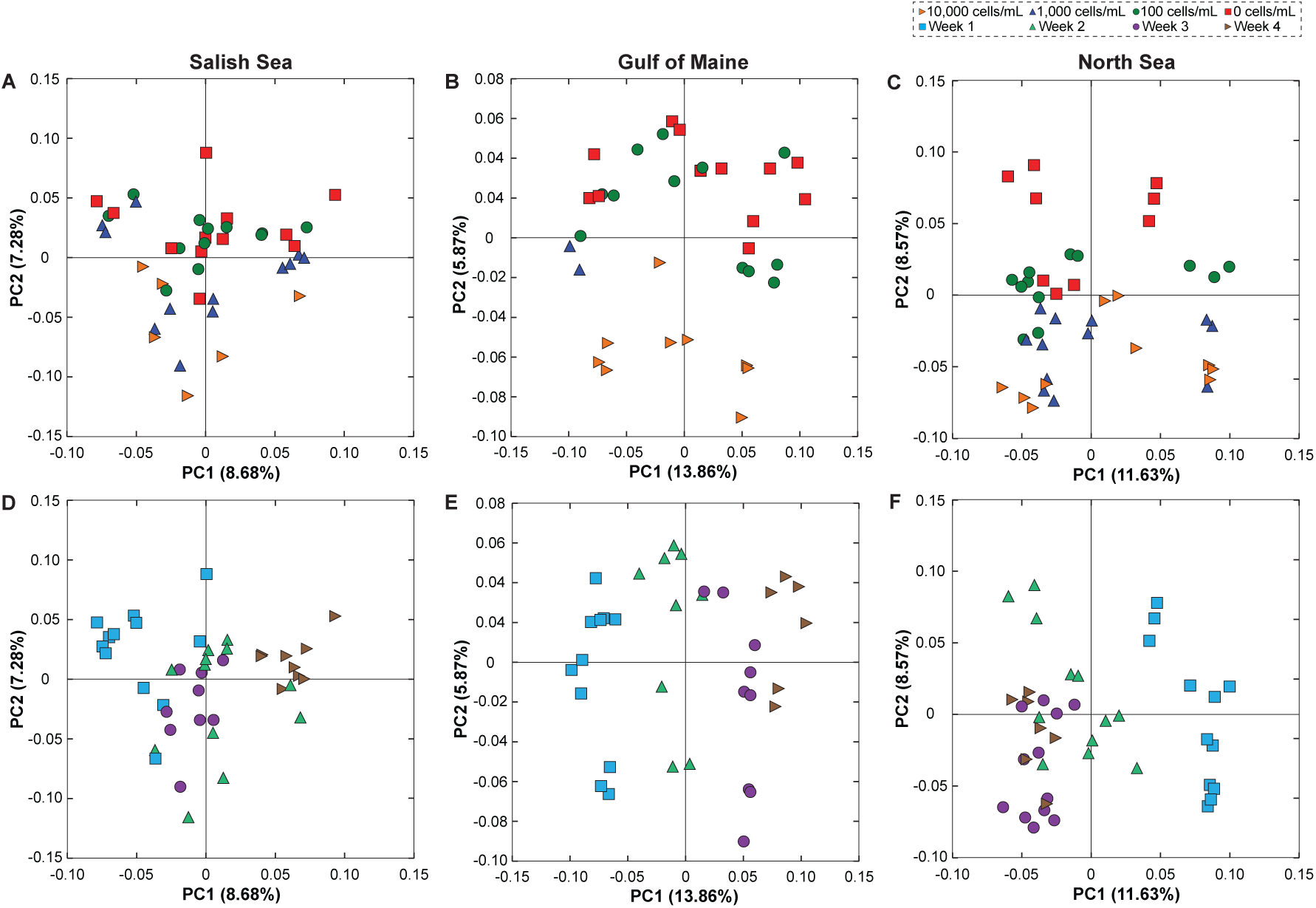
Dietary and temporal shifts in the bacterial communities associated with *Strongylocentrotus droebachiensis* larvae. Community similarity based on *S. droebachiensis* larval-associated bacterial communities based on food availability (A-C) over a multi-week exposure (D-F) for populations from the Salish Sea (A, D), Gulf of Maine (B, E), and North Sea (C, F). Comparisons between food availability (A-C) and over time (D-F) are based on unweighted UniFrac comparisons. (Note: this is the unweighted complement to Fig. 2.)

**Figure S5.**
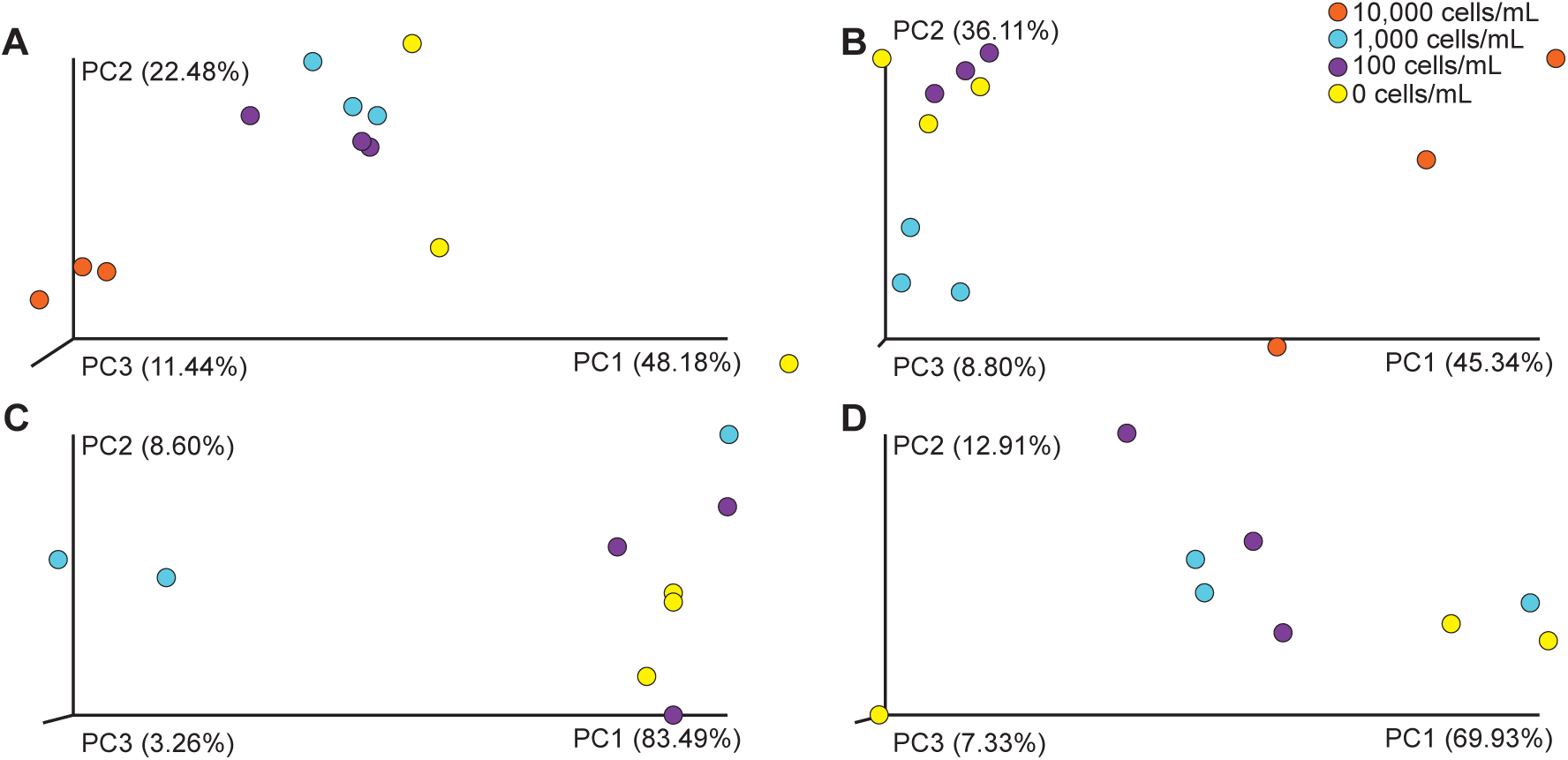
Community similarity of *Strongylocentrotus droebachiensis* larvae from the Salish Sea across time. Similarity of bacterial communities associated with *S. droebachiensis* larvae from the Salish Sea for one (A; p < 0.002), two (B; p < 0.001), three (C; p < 0.001), or four (D; p < 0.001) weeks and having been fed 10,000 (orange), 1,000 (blue), 100 (purple), or 0 (yellow) cells•mL^−1^.

**Figure S6.**
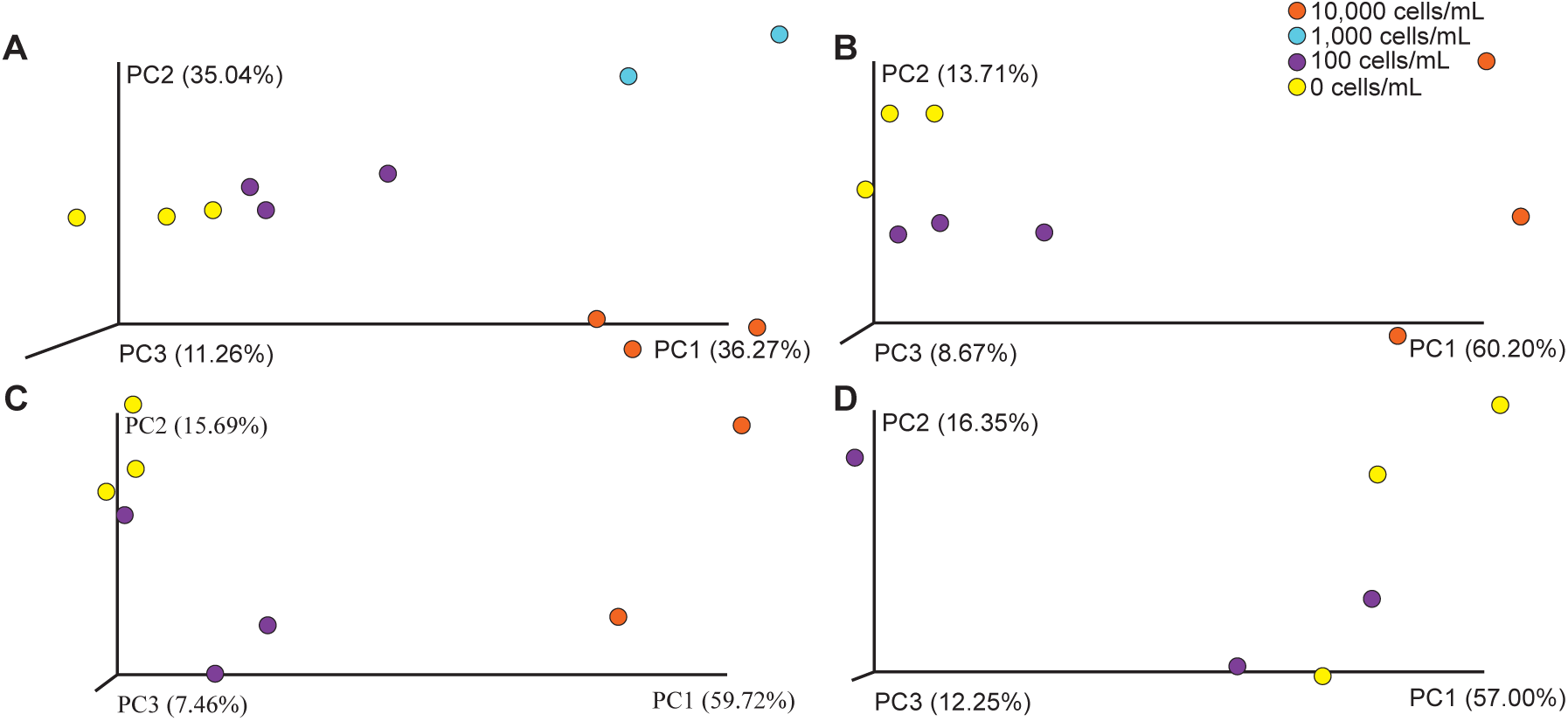
Community similarity of *Strongylocentrotus droebachiensis* larvae from the Gulf of Maine across time. Similarity of bacterial communities associated with *S. droebachiensis* larvae from the Gulf of Maine for one (A; p < 0.001), two (B; p < 0.005), three (C; p < 0.007), or four (D; p = 0.430) weeks and having been fed 10,000 (orange), 1,000 (blue), 100 (purple), or 0 (yellow) cells•mL^−1^.

**Figure S7.**
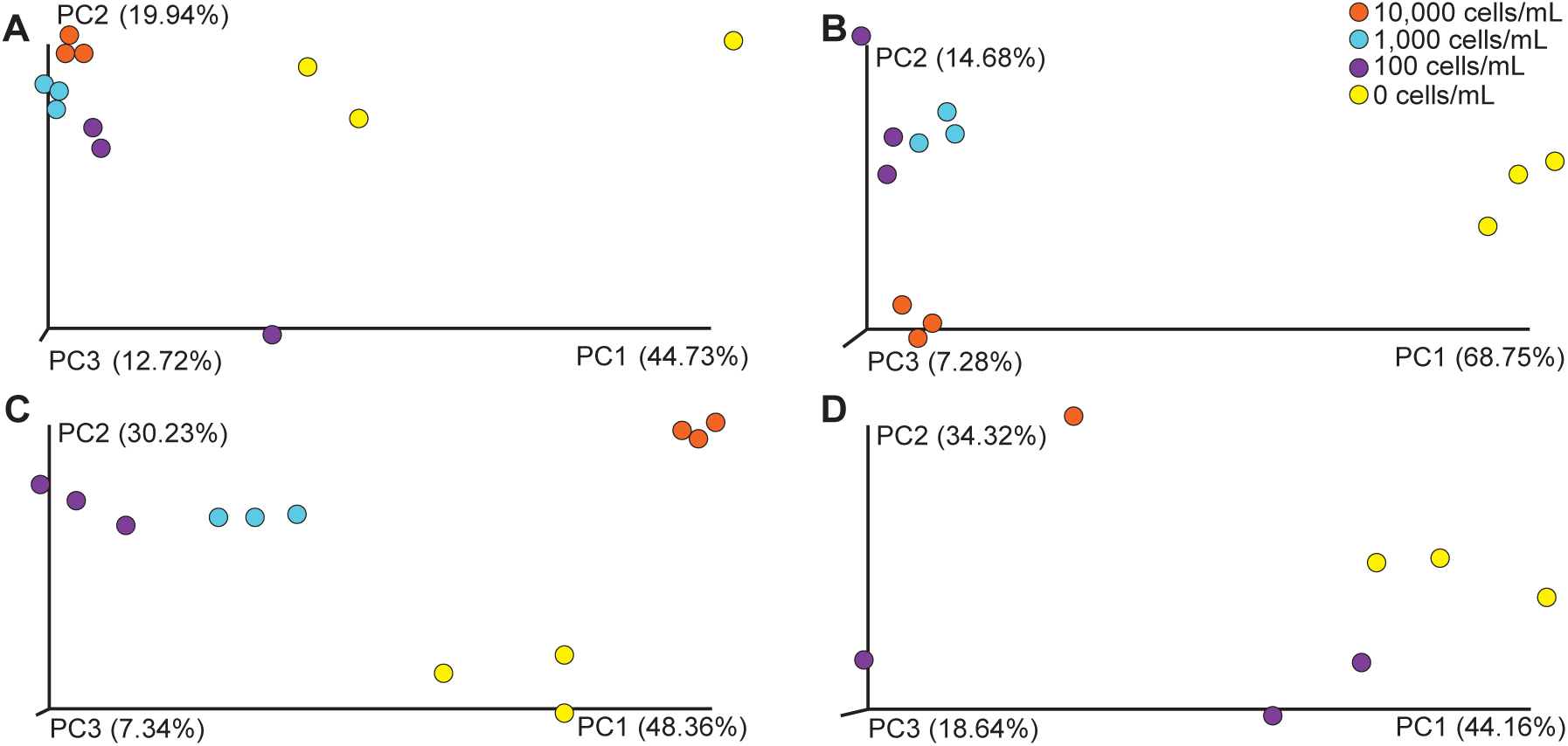
Community similarity of *Strongylocentrotus droebachiensis* larvae from the North Sea across time. Similarity of bacterial communities associated with *S. droebachiensis* larvae from the North Sea for one (A; p < 0.002), two (B; p < 0.001), three (C; p < 0.001), or four (D; p < 0.020) weeks and having been fed 10,000 (orange), 1,000 (blue), 100 (purple), or 0 (yellow) cells•mL^−1^.

**Figure S8.**
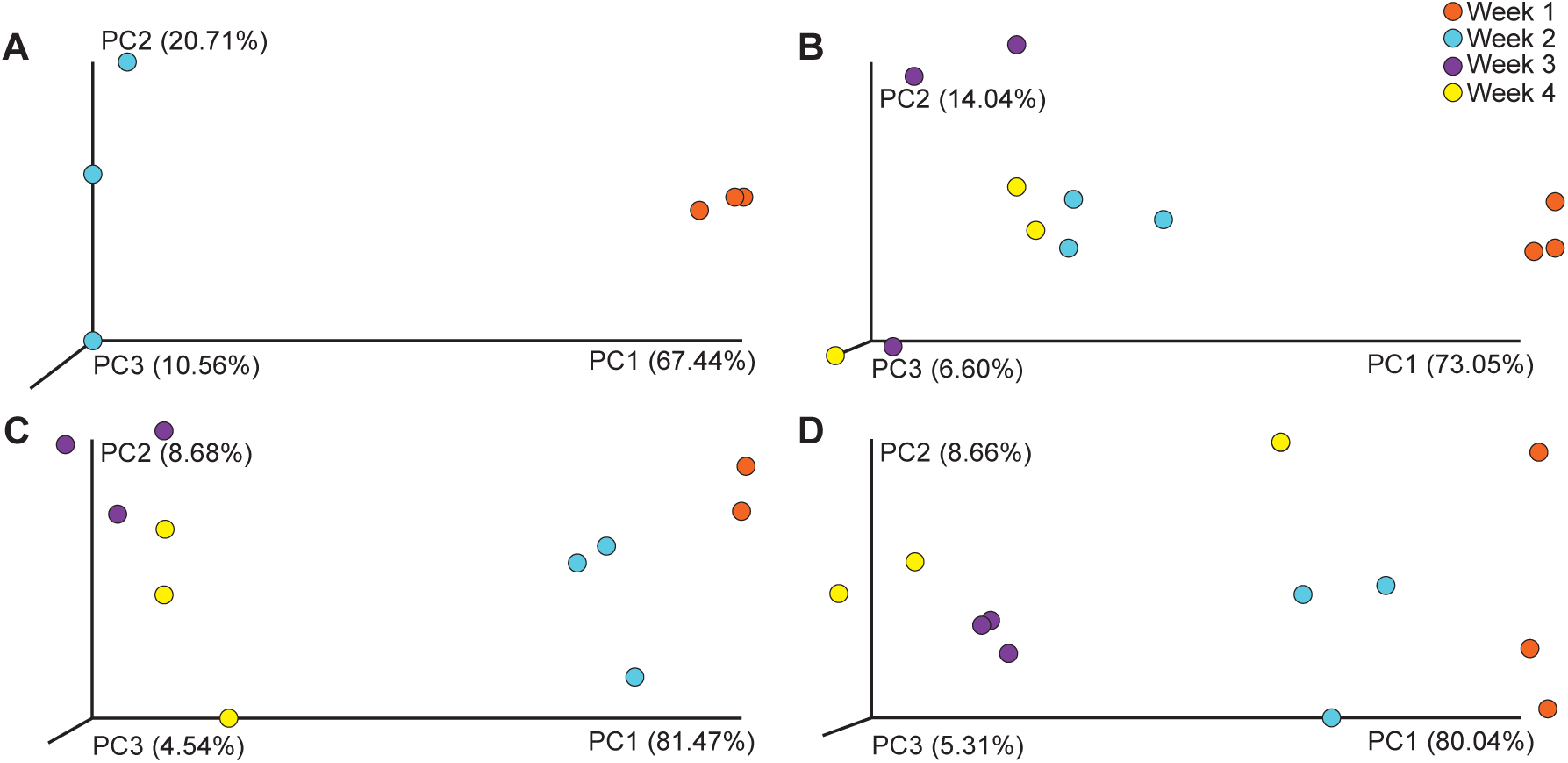
Community similarity of *Strongylocentrotus droebachiensis* larvae from the Salish Sea across diets. Similarity of bacterial communities associated with *S. droebachiensis* larvae from the Salish Sea having been fed 10,000 (A; p < 0.001), 1,000 (B; p < 0.001), 100 (C; p < 0.001), or 0 (D; p < 0.001) cells•mL^−1^ for one (orange), two (blue), three (purple), or four (yellow) weeks.

**Figure S9.**
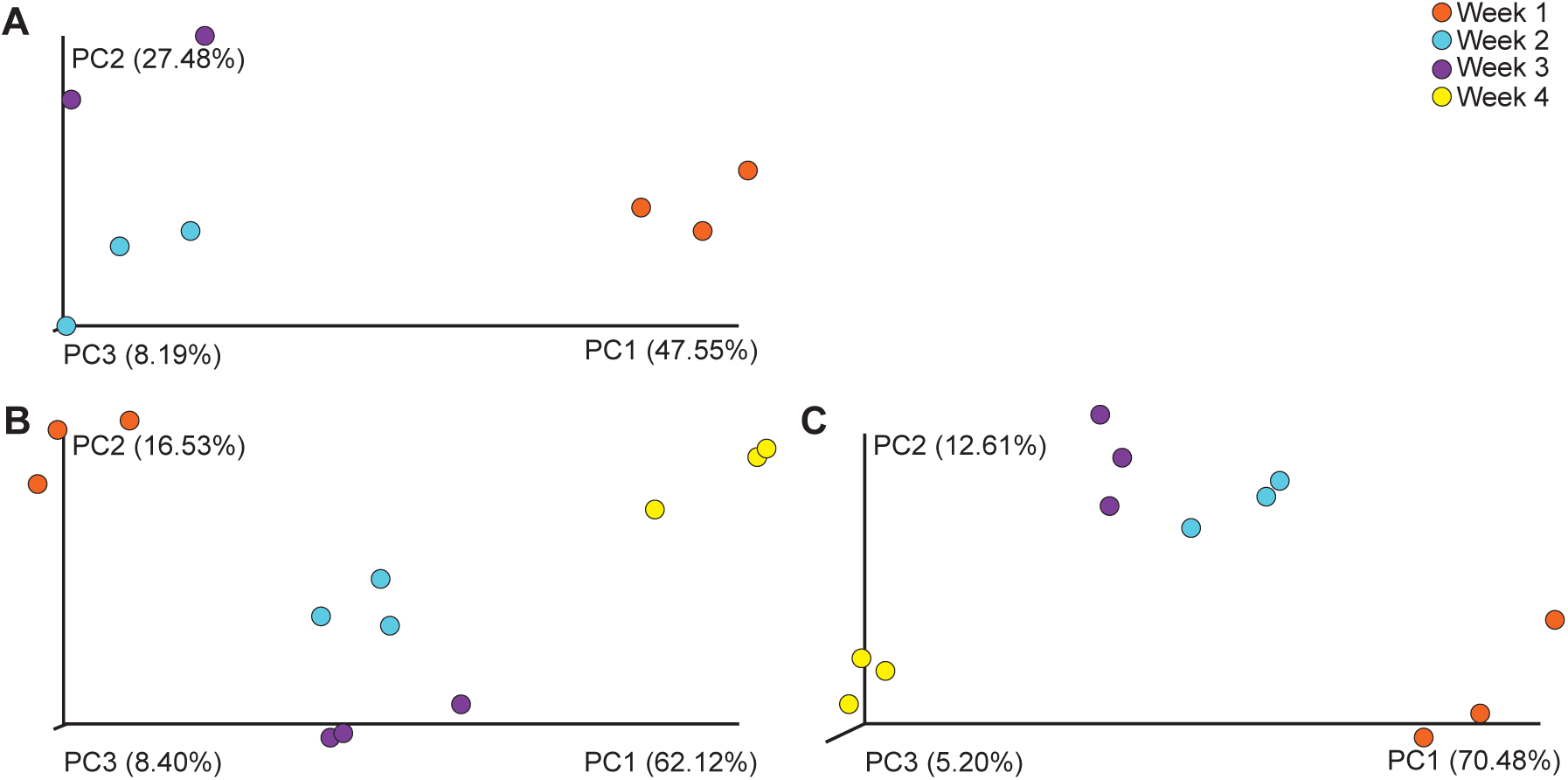
Community similarity of *Strongylocentrotus droebachiensis* larvae from the Gulf of Maine across diets. Similarity of bacterial communities associated with *S. droebachiensis* larvae from the Gulf of Maine having been fed 10,000 (A; p < 0.005), 100 (B; p < 0.001), or 0 (C; p < 0.002) cells•mL^−1^ for one (orange), two (blue), three (purple), or four (yellow) weeks.

**Figure S10.**
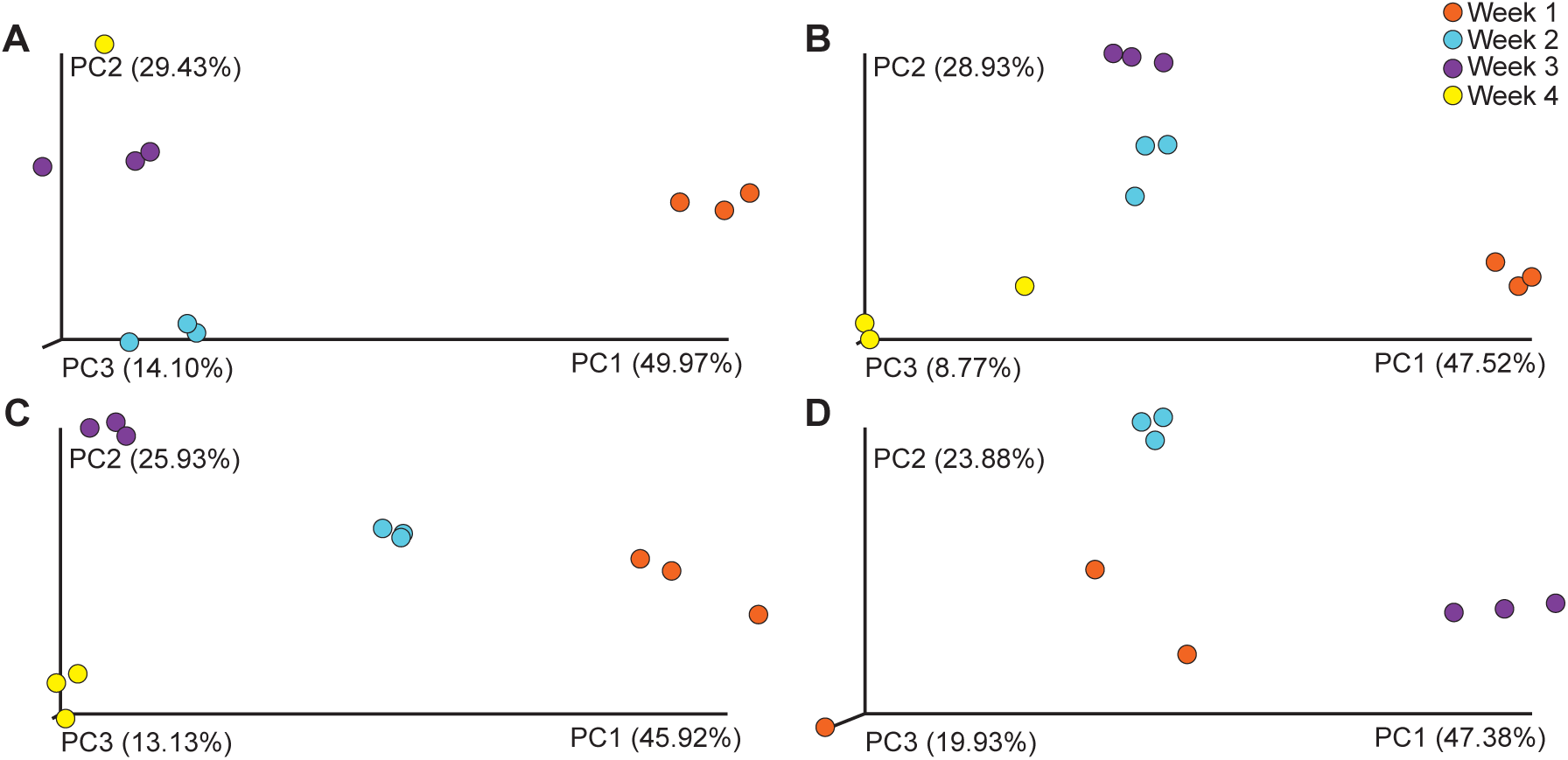
Community similarity of *Strongylocentrotus droebachiensis* larvae from the North Sea across diets. Similarity of bacterial communities associated with *S. droebachiensis* larvae from the North Sea having been fed 10,000 (A; p < 0.002), 1,000 (B; p < 0.002), 100 (C; p < 0.002), or 0 (D; p < 0.006) cells•mL^−1^ for one (orange), two (blue), three (purple), or four (yellow) weeks.

**Figure S11.**
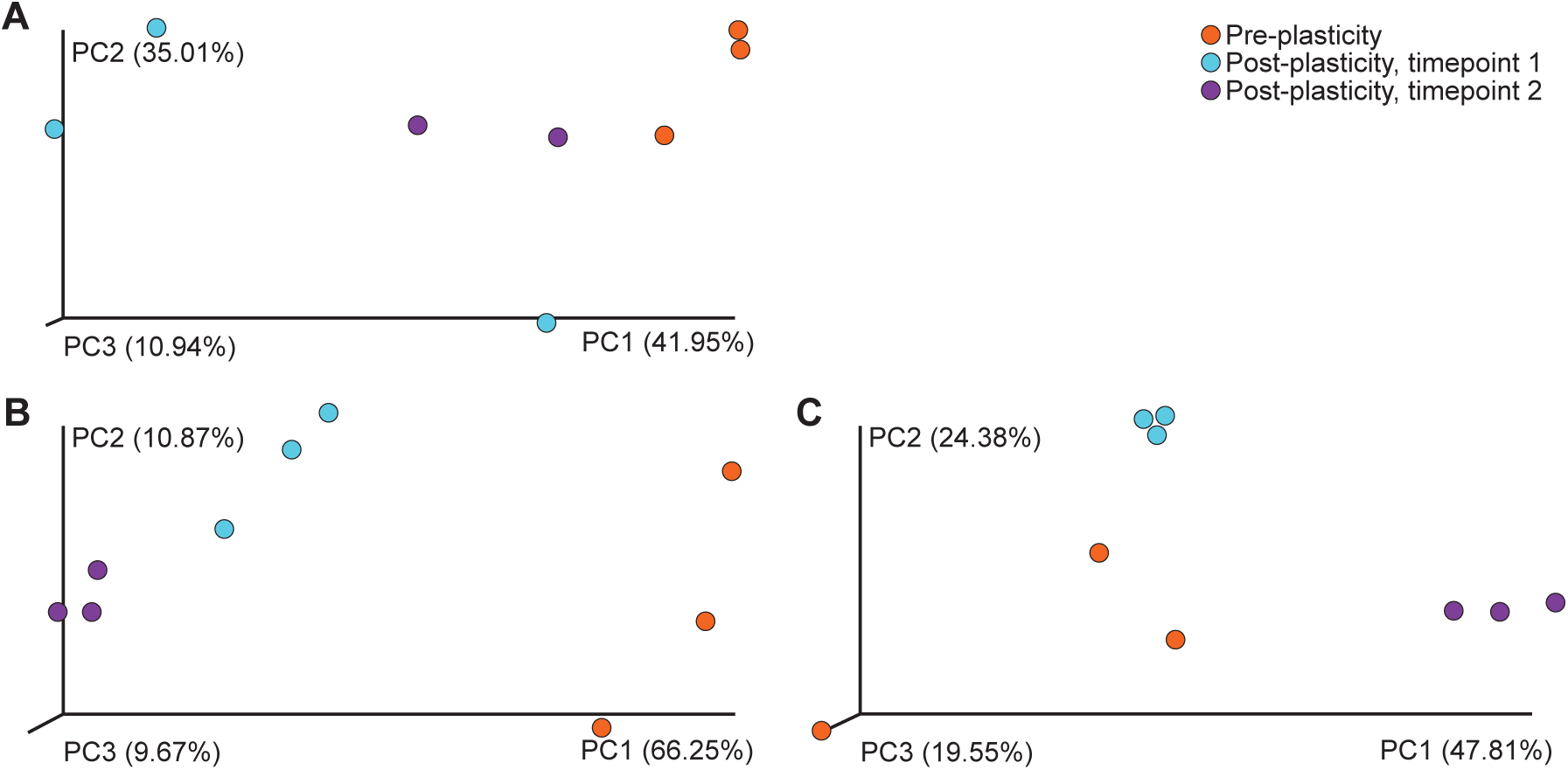
Community similarity of the associated microbial community along a phenotypic continuum for *Strongylocentrotus droebachiensis* from three geographic locations. Similarity of bacterial communities associated with *S. droebachiensis* larvae from the Salish Sea (A; p = 0.046), Gulf of Maine (B; p < 0.005), and North Sea (C; p < 0.004) pre (orange) and post (blue and purple) expression of plasticity.

**Figure S12.**
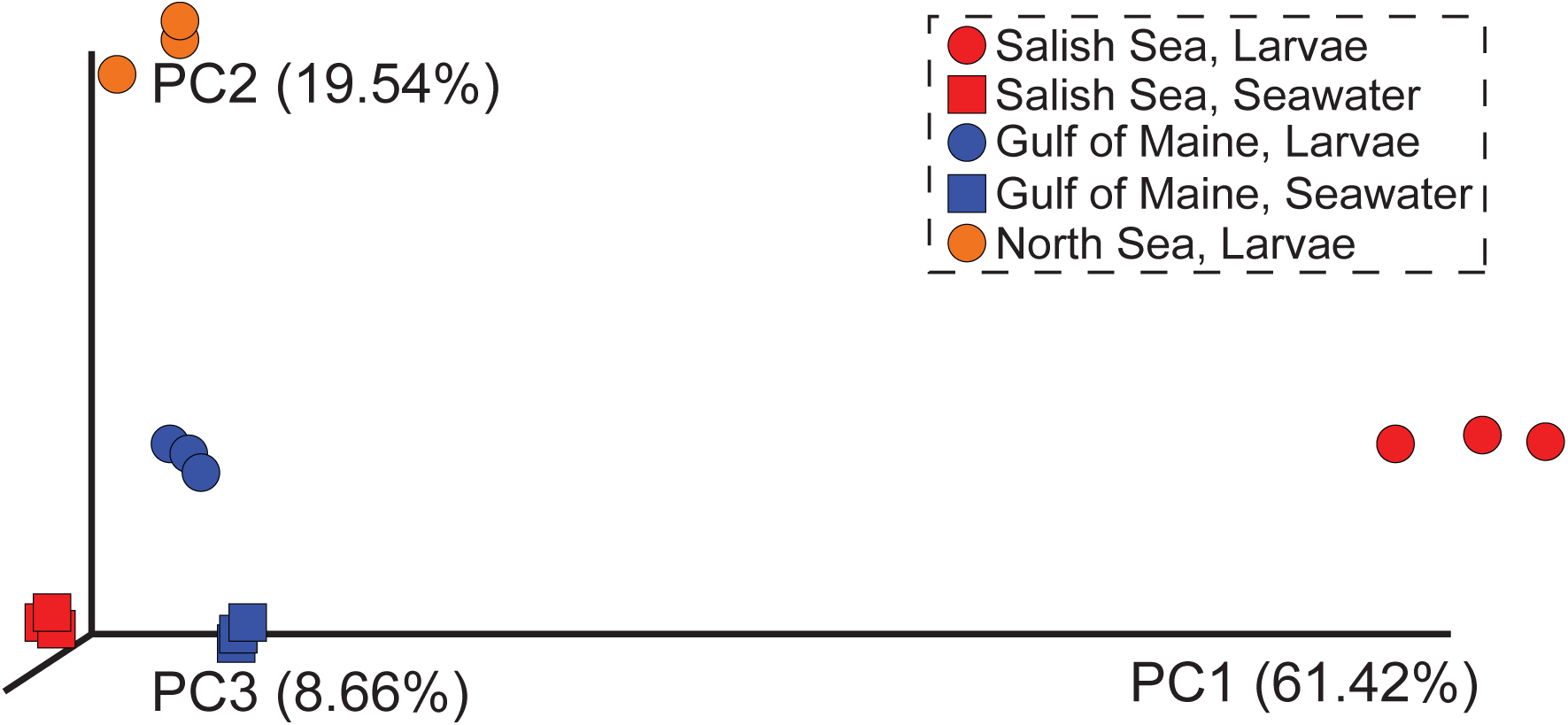
Similarity of the associated microbial community for *Strongylocentrotus droebachiensis* larvae from three geographic locations. Community similarity (p < 0.001) of the associated microbiota between *S. droebachiensis* larvae (circles) from the Salish Sea (blue), Gulf of Maine (red), and North Sea (yellow) and seawater (squares).

**Figure S13.**
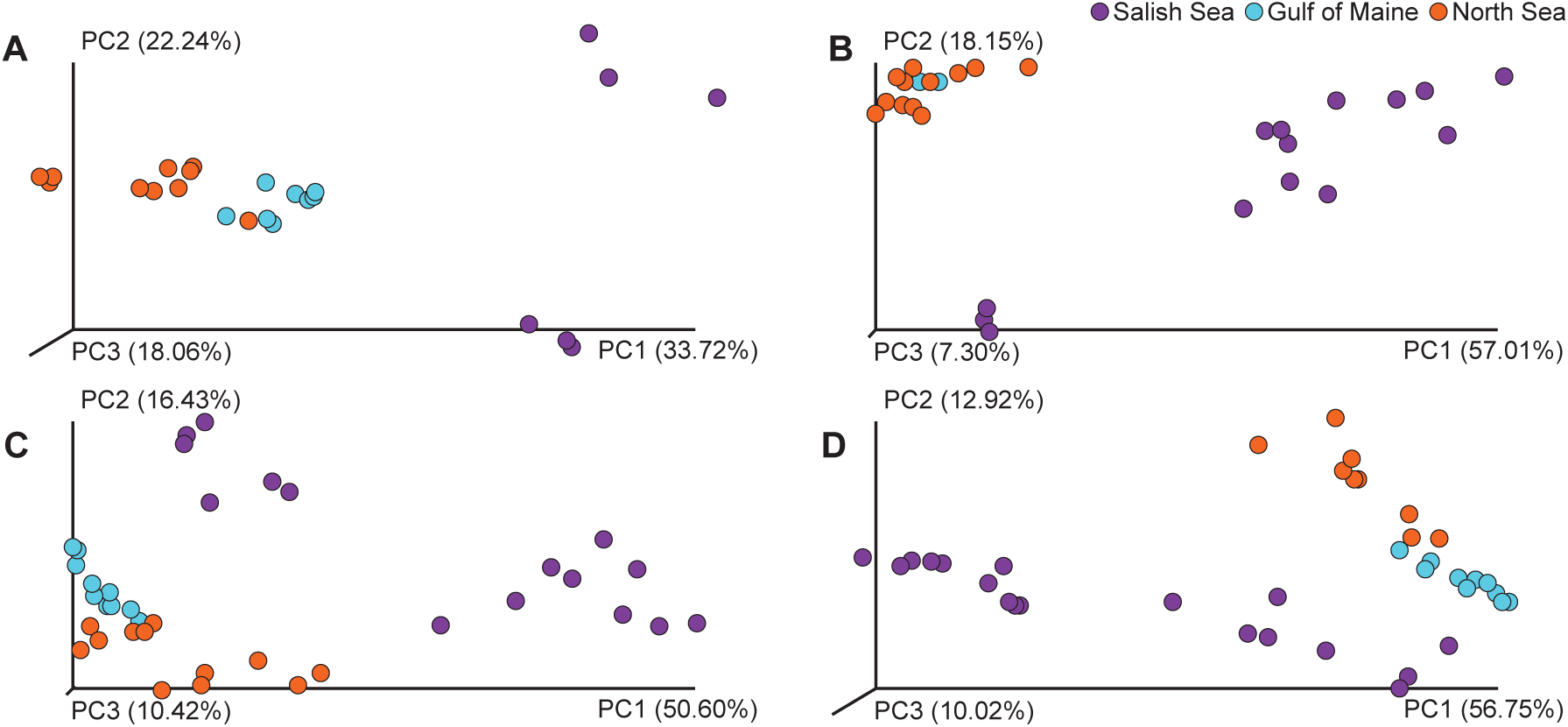
Community similarity of *Strongylocentrotus droebachiensis* larvae across diets. Similarity of bacterial communities associated with *S. droebachiensis* larvae from the Salish Sea (purple), Gulf of Maine (blue), or North Sea (orange) and having been fed 10,000 (A; p < 0.001), 1,000 (B; p < 0.001), 100 (C; p < 0.001), and 0 (D; p < 0.001) cells•mL^−1^.

**Figure S14.**
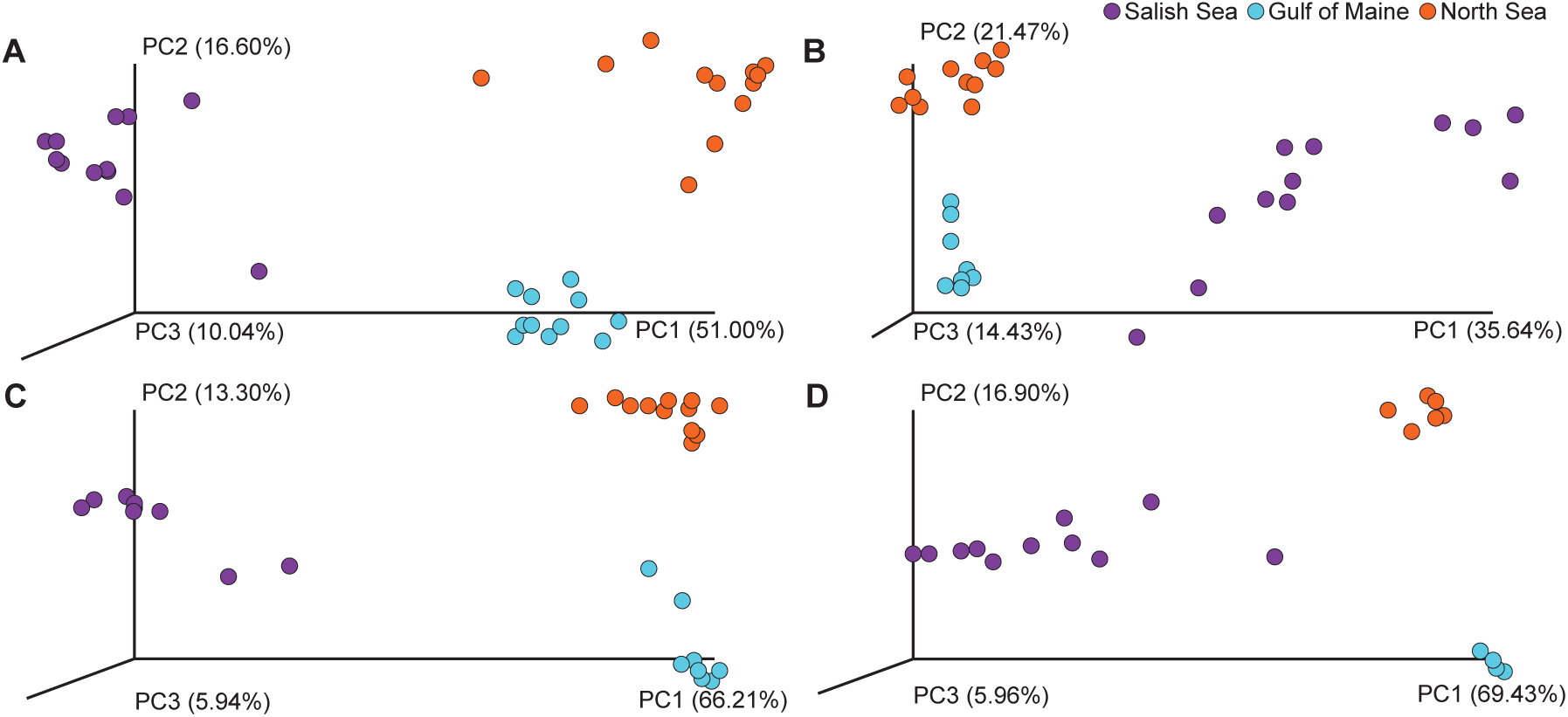
Community similarity of *Strongylocentrotus droebachiensis* larvae across time. Similarity of bacterial communities associated with *S. droebachiensis* larvae from the Salish Sea (purple), Gulf of Maine (blue), or North Sea (orange) and having been differentially fed for one (A; p < 0.001), two (B; p < 0.001), three (C; p < 0.001), or four (D; p < 0.001) weeks.

## Note S1

**QIIME analysis**

Starting with raw read files from MiSeq:

1) Pair forward and reverse files using PEAR (pear-0.9.10-bin-64)

2) Trim paired files using Trimmomatic (trimmomatic-0.36.jar)

3) Convert paired and trimmed .fastq to .fasta using the custom code: cat [input .fastq] | paste – - - - | cut -f 1,2 | sed 's/^/>/'| tr "\t" "\n" >[output .fasta]

4) Validate tab-delimited mapping file using "validate_mapping_file.py"

5) Generate meta-.fasta files using "add_qiime_labels.py"

6) Detect chimeras from meta-.fasta (called, "combined_seqs.fna")

7) Filter chimeras using "filter_fasta.py"

8) Pick OTUs using "pick_open_reference_otus.py"

9) Filter OTUs with >10 reads using "filter_otus_from_otu_table.py"

10) Filter ‘o_Cryptophyta’ using "filter_taxa_from_otu_table.py"

11) Determine rarefaction depth using "biom summarize-table"

12) Filtered .biom table was rarified using “single_rarefaction.py”

13) Filtered filtered .biom table was split using "split_otu_table.py" to test specific hypotheses

14) Alpha diversity was compared using “alpha_diversity.py”

15) Beta diversity was compared using "jackknifed_beta_diversity.py" and 2D PCoA plots of unweighted and weighted UniFrac values were generated using “make_2d_plots.py”

16) Statistical comparisons of unweighted and weighted UniFrac were performed with “compare_categories.py” using the ‘--method’ “anosim,” “adonis,” and “permanova.”

17) Taxonomic plots were generated using "summarize_taxa_through_plots.py"

18) Unique and shared OTUs were determined both using “compute_core_microbiome.py”and “shared_phylotypes.py”

